# Unveiling the Immune Landscape of COVID-19 and prolonged Long-COVID through Single-Cell RNA Sequencing

**DOI:** 10.1101/2025.03.05.641671

**Authors:** Marta Līva Spriņģe, Kristīne Vaivode, Rihards Saksis, Nineļa Miriama Vainšeļbauma, Laura Ansone, Monta Brīvība, Helvijs Niedra, Vita Rovīte

## Abstract

Long-COVID affects at least 10% of COVID-19 survivors, displaying debilitating symptoms across multiple organ systems. Despite the increasing prevalence, the underlying causes remain unclear. This study presents a unique analysis of the PBMC transcriptomic landscape of COVID-19 and Long-COVID patients at a single-cell resolution. We reconstructed the cell state and communication using differentially expressed gene profiling and ligand-receptor interaction analyses. Our results reveal altered T and NK cell subset proportions, diminished proliferating lymphocyte and B cell signalling capacity, and the expression of exhaustion and cytotoxicity associated genes 1.5 – 2 years post-infection, suggesting incomplete immune recovery. Collectively, these findings provide insights into the immune processes underlying the progression of COVID-19 into a chronic Long-COVID state.

## Introduction

The SARS-CoV-2 virus (COVID-19) emerged in late 2019, and soon after, on the 11th of March 2020, this pathogen was announced to have become the cause of a global pandemic (World Health Organisation). Tragically causing over 7 million deaths and affecting the lives of millions of patients to date (World Health Organization, 2025), the virus can introduce a plethora of symptoms, including chest pain, shortness of breath, and cognitive dysfunction, that may persist or develop even after initial COVID-19 infection has been resolved (Al-Aly *et al*, 2021; Davis *et al*, 2023). Commonly referred to as Long-COVID, other umbrella terms such as “Post-COVID-19 condition”, “Post-acute COVID syndrome”, and “Post-acute sequelae of SARS-CoV-2” are often used interchangeably to describe the various symptoms persisting for at least three months after acute COVID-19 infection, with no other underlying cause (World Health Organisation, 2022).

It has been generally accepted that Long-COVID develops in 10-20% of COVID-19 survivors (World Health Organisation, 2022). However, recent studies show the general incidence of Long-COVID may have been underestimated, disproportionately affecting different groups of the population (Bucciarelli *et al*, 2022; Davis *et al*, 2021; Hastie *et al*, 2023; Kim *et al*, 2023; Mantovani *et al*, 2022; Han *et al*, 2022). Several studies show women are at a higher risk of developing Long-COVID (Bucciarelli *et al*, 2022; Han *et al*, 2022). Severe acute COVID-19 also considerably increases the likelihood of developing Long-COVID (Han *et al*, 2022). Smoking habits, age, and diabetes mellitus have been cited as risk factors as well (Sugiyama *et al*, 2024). Conflicting evidence has emerged regarding the association of Long-COVID with vaccination history (Abu Hamdh & Nazzal, 2023; Kim *et al*, 2023; Romero-Ibarguengoitia *et al*, 2024) or virus variants (Sugiyama *et al*, 2024; Davis *et al*, 2023), which remains to be studied.

Long-COVID, with its symptom spectrum ranging from mild to severe, can become profoundly debilitating for many patients affected by the condition (Malik *et al*, 2022). Acute COVID-19, as well as Long-COVID, manifests across multiple organ systems; notably, pulmonary and cardiovascular complications pose a serious threat to the life and health span of the patient (Al-Aly *et al*, 2021; Xie *et al*, 2022; Davis *et al*, 2023). Beyond the known incidence of patients presenting respiratory distress and cardiovascular events during the acute phase of COVID-19, an increase in a range of associated complications in the cases that develop Long-COVID, including myocarditis, dyspnoea, and the requirement of oxygen support has also been observed (Al-Aly *et al*, 2021; Xie *et al*, 2022; Davis *et al*, 2023). Research points to immune system dysfunction during COVID-19, which may extend to prolonged dysregulation and contribute to lasting complications in Long-COVID cases (Yin et al., 2024).

Characterizing the dynamic response and functional alterations of T cells in COVID-19 has been essential to understanding the virus-host immune system interactions (Alahdal & Elkord, 2022; Rha & Shin, 2021). Recently, specific subsets of CD8^+^ T cells expressing *PD1* and *CTLA4* were found in Long-COVID cases 8 months post-acute infection (Yin *et al*, 2024). These markers are associated with T cell exhaustion, which is driven by chronic antigen stimulation and is known to play a role in persistent viral infections (Alahdal & Elkord, 2022; Rha & Shin, 2021). Furthermore, severe COVID-19 has been shown to induce lasting effects on the function of the innate immune system. Monocyte populations in Long-COVID peripheral blood mononuclear cells (PBMCs) have been described as dysfunctional (Scott *et al*, 2023), hyper-responsive (Cheong *et al*, 2023), or altered in other ways (Junqueira *et al*, 2022). However, their precise role in driving Long-COVID pathogenesis and severe COVID-19 states remains unclear. Several different immune mechanisms underlying lasting COVID-19 complications have been studied, yet whether these are involved in promoting, sustaining, or aggravating Long-COVID remains to be elucidated. Building further on the studies mentioned above and our previous work (Vaivode *et al*, 2024), where we observe increased immature neutrophil counts in a Long-COVID patient PBMCs 3 months post-infection, along with a large mature neutrophil population at 2 years post-infection, we aim to investigate these immune alterations temporally in a larger patient cohort.

Given the sudden onset and rapid spread of the SARS-CoV-2 virus, it is crucial to study the long-term effects and implications on patient life and health span this infection poses. Long-COVID cases are rising, and the number of patients requiring long-term post-COVID-19 care is increasing (Davis *et al*, 2023). It is crucial to follow up with patients, study the symptoms presenting after acute infection resolution, and inquire deeper into researching the mechanisms guiding systemic Long-COVID conditions.

To further investigate the involvement of immune cells in acute and Long-COVID-19, we use single-cell RNA sequencing (scRNA-seq) for the characterization of PBMCs from a select cohort of patients during acute COVID-19 infection, 3 months and 1.5 – 2 years post-acute infection. We obtained blood samples from female patients who had been hospitalized during acute infection and established three study groups: those who later developed Long-COVID with lasting cardiovascular (hereinafter referred to as LC-CV) and those developing pulmonary complications (LC-Pulm), along with samples from patients who did not develop Long-COVID (Non-LC). Specifically, we focus on female patients with severe initial infection, as gender and disease severity have been recognized as factors for an increased risk for developing Long-COVID (Bucciarelli *et al*, 2022; Han *et al*, 2022). ScRNA-seq allows the identification of individual cells, classification of subsets, and analysis of cellular interactions involved, which may lead to possible answers regarding the pathophysiological principles of Long-COVID. In the present study, we aim to unravel the intricate details of immune cell transcriptomic alterations and the immune system perturbations involved in acute and chronic COVID-19 conditions in a longitudinal manner.

## Results

To investigate the immune cell landscape of patients with acute COVID-19 infection, those who had fully recovered or developed Long-COVID (LC), implemented scRNA-seq for PBMC analysis. We obtained RNA transcripts at a single-cell resolution and filtered the high-quality cells, resulting in a total of 174 336 cells from 9 patients. Each patient included in the study was classified as either having recovered (Non-LC, n = 3) or developing cardiovascular (LC-CV, n = 3) or pulmonary (LC-Pulm, n = 3) Long-COVID complications. The study design is depicted in Fig. 1A. All patients were hospitalized during the acute phase of COVID-19 infection. Blood samples were obtained from these patients during the acute phase, after 3 months, and 1.5 – 2 years post-infection (Fig. 1A).

**Figure 1.**
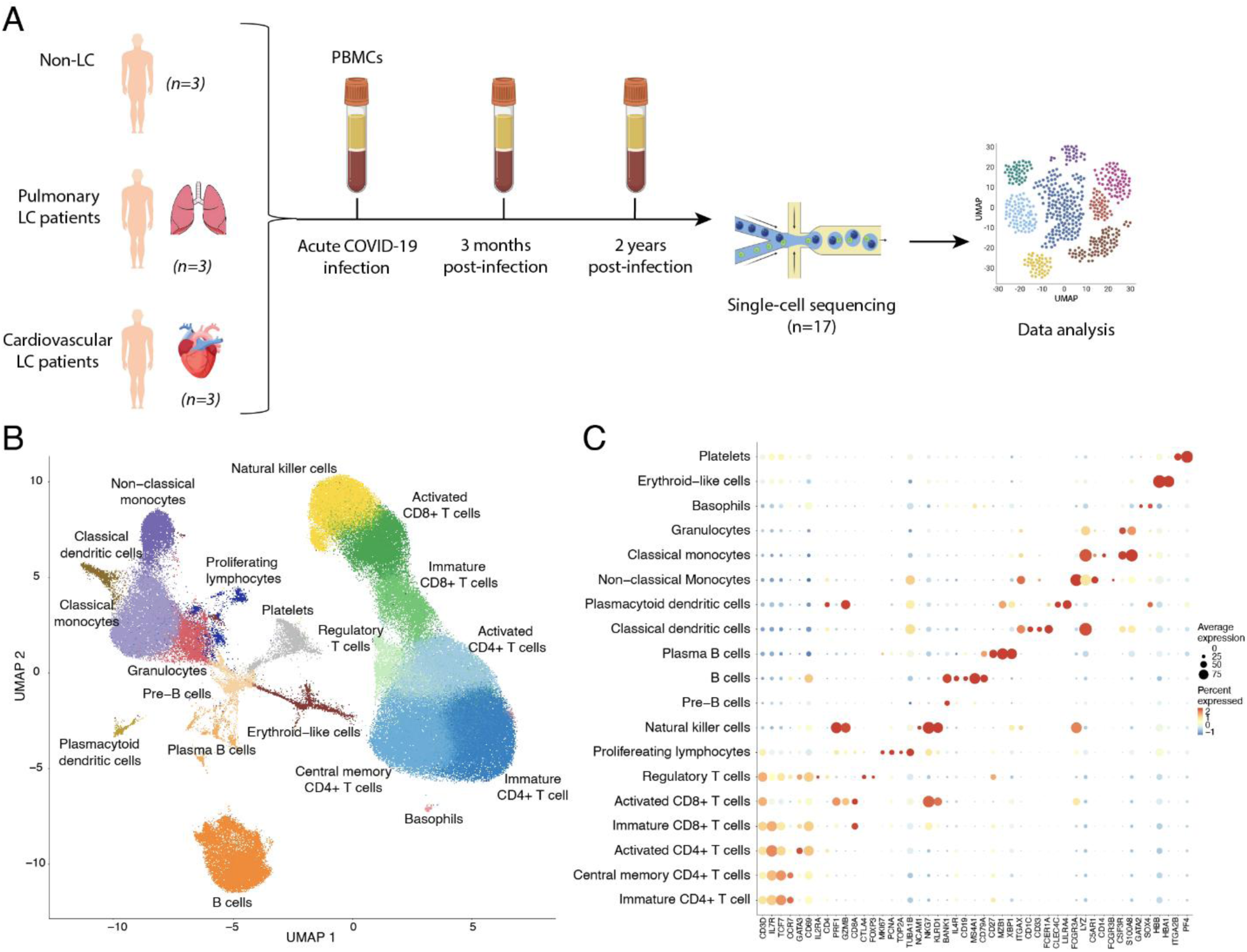
Study design and cell population identification. A Overview of the study design. The workflow consists of PBMC isolation using blood samples from 9 patients, single-cell RNA sequencing, and data analysis. Samples were obtained from patients during the acute phase of infection, after 3 months, and after 1.5 – 2 years. Patients who developed cardiovascular (LC-CV, n = 3) and pulmonary (LC-Pulm, n = 3) Long-COVID complications were selected for the study. Patients who did not develop Long-COVID (Non-LC, n = 3) were used as controls. B Integrated single-cell population landscape across all conditions and samples. The UMAP projection shows a total of 174 336 cells defined in 22 clusters and 19 cell populations. Each point represents a single cell coloured according to the inferred cell type. Cell type identification based on ScType (Ianevski *et al*, 2022) and gene marker average expression and prevalence, as shown in C. C Gene expression markers used to differentiate cell populations and subpopulations. The dot plot shows the identified cell populations along the y-axis and used marker genes along the x-axis. Dot size indicates the average expression of the marker gene in each of the identified populations. The dot colour indicates the percentage of cells in each population in which the marker gene was expressed.

Results of the cell analysis were presented on a UMAP graph showing 22 cell clusters, representing proximity in gene expression across individual cells (Fig. 1B). Clusters were identified as cell types according to ScType (Ianevski *et al*, 2022) and well-established marker genes (Tab. EV1), as shown in Fig. 1C. A total of 19 cell-type clusters were identified representing blood immune cells (Tab. EV1, Fig. 1C).

**Expanded View Table 1.**
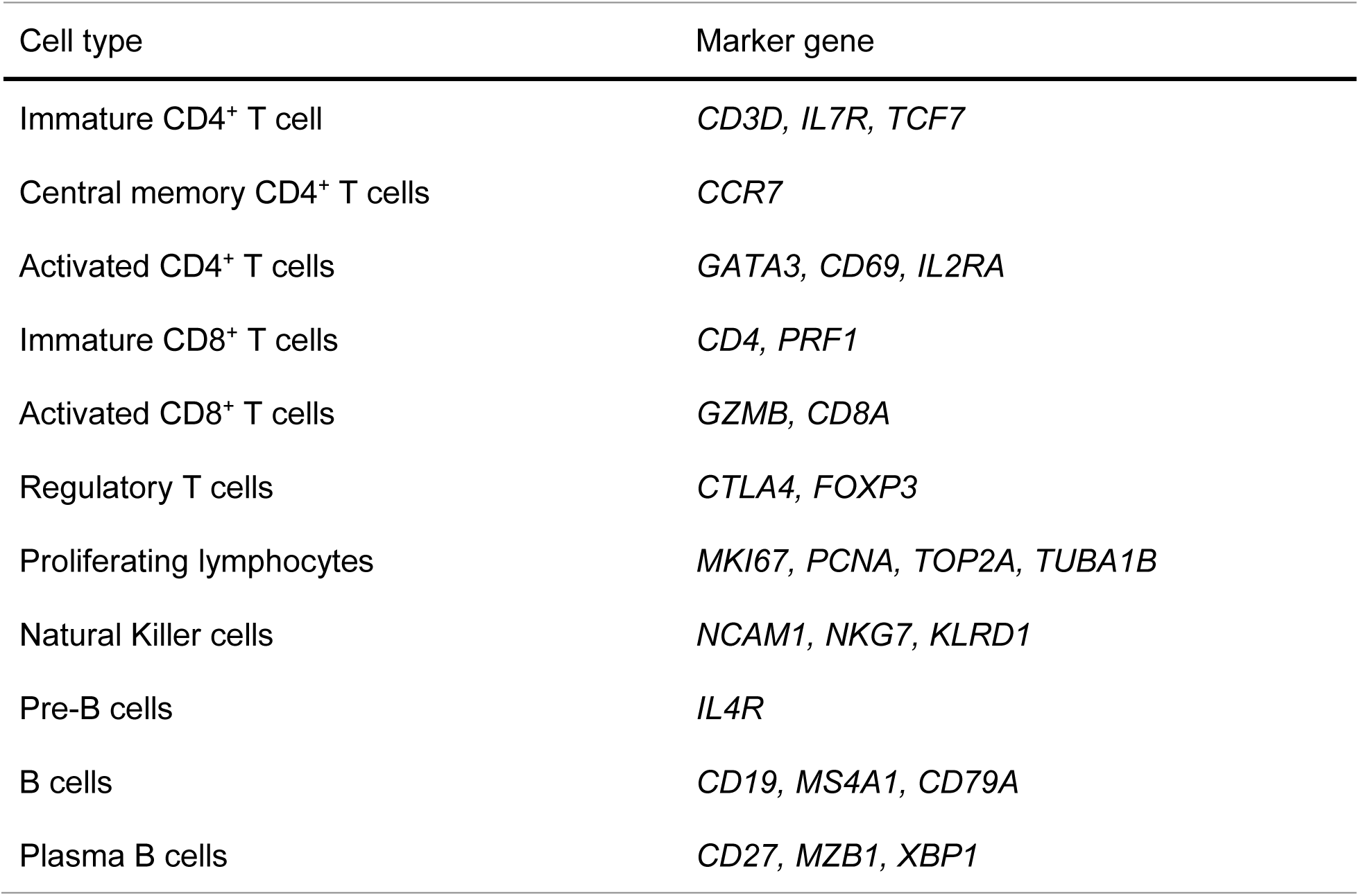
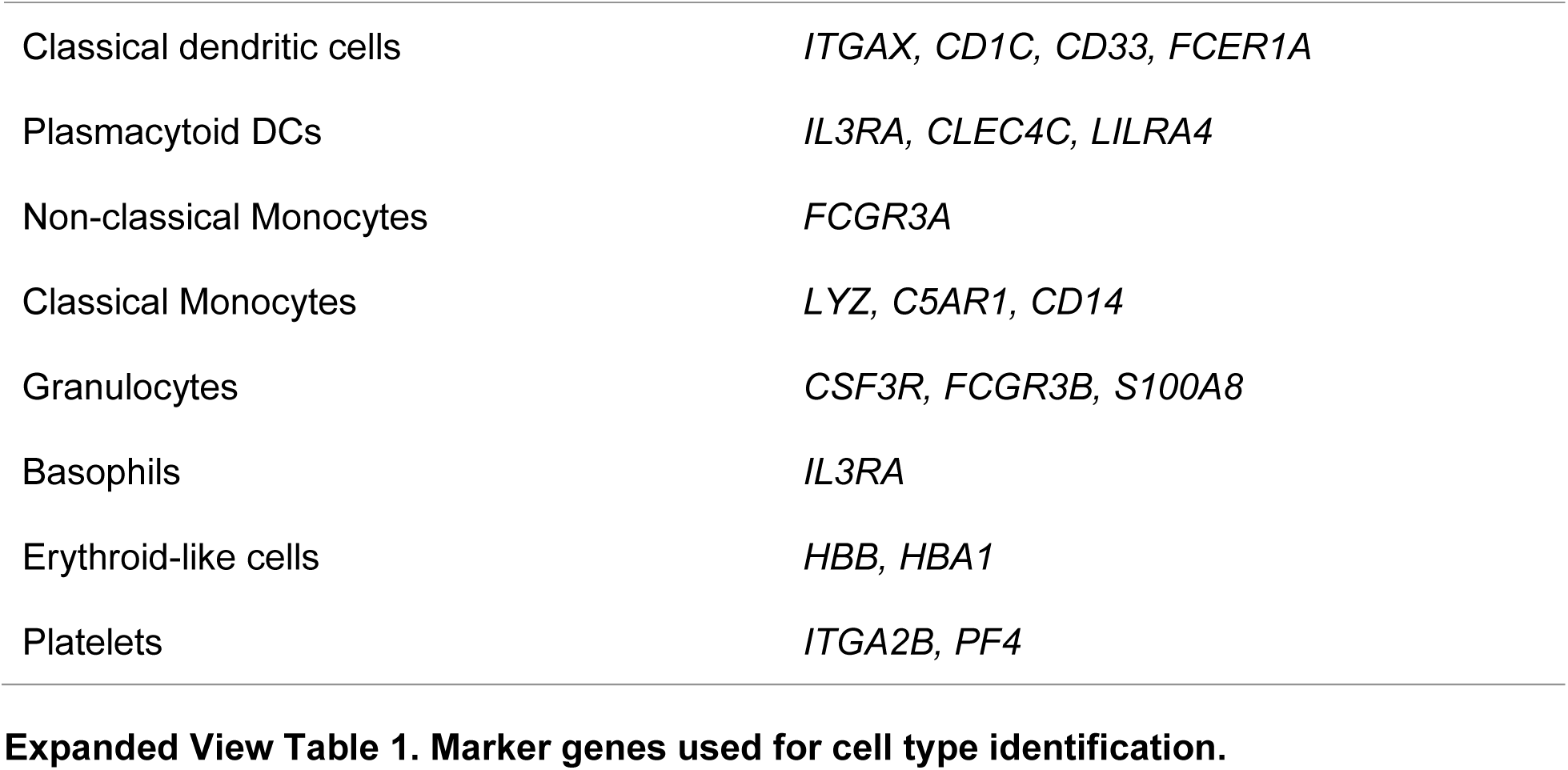
Marker genes used for cell type identification.

| Cell type | Marker gene |
| --- | --- |
| Immature CD4<sup>+</sup> T cell | *CD3D, IL7R, TCF7* |
| Central memory CD4<sup>+</sup> T cells | *CCR7* |
| Activated CD4<sup>+</sup> T cells | *GATA3, CD69, IL2RA* |
| Immature CD8<sup>+</sup> T cells | *CD4, PRF1* |
| Activated CD8<sup>+</sup> T cells | *GZMB, CD8A* |
| Regulatory T cells | *CTLA4, FOXP3* |
| Proliferating lymphocytes | *MKI67, PCNA, TOP2A, TUBA1B* |
| Natural Killer cells | *NCAM1, NKG7, KLRD1* |
| Pre-B cells | *IL4R* |
| B cells | *CD19, MS4A1, CD79A* |
| Plasma B cells | *CD27, MZB1, XBP1* |
| Classical dendritic cells | *ITGAX, CD1C, CD33, FCER1A* |
| Plasmacytoid DCs | *IL3RA, CLEC4C, LILRA4* |
| Non-classical Monocytes | *FCGR3A* |
| Classical Monocytes | *LYZ, C5AR1, CD14* |
| Granulocytes | *CSF3R, FCGR3B, S100A8* |
| Basophils | *IL3RA* |
| Erythroid-like cells | *HBB, HBA1* |
| Platelets | *ITGA2B, PF4* |

We further examined the differences in immune cell populations between patient groups based on the identified clusters. The UMAP overlay of the immune cell landscapes of Non-LC, LC- Pulm and LC-CV complication groups show the differential cell clustering between disease conditions (Fig. 2A). Of note, proliferative lymphocytes displayed unique clustering patterns between the three groups. While distinct clustering can be observed between conditions throughout the visualization, B cell subpopulations, as well as platelets, non-classical monocytes (Mono), plasmacytoid dendritic cells (DCs), and T cell subpopulations diverge most considerably (Fig. 2A). The patient cell landscape between the three timepoints (acute phase, 3 months and 1.5 – 2 years post-infection) is depicted in Appendix Fig. S1.

**Figure 2.**
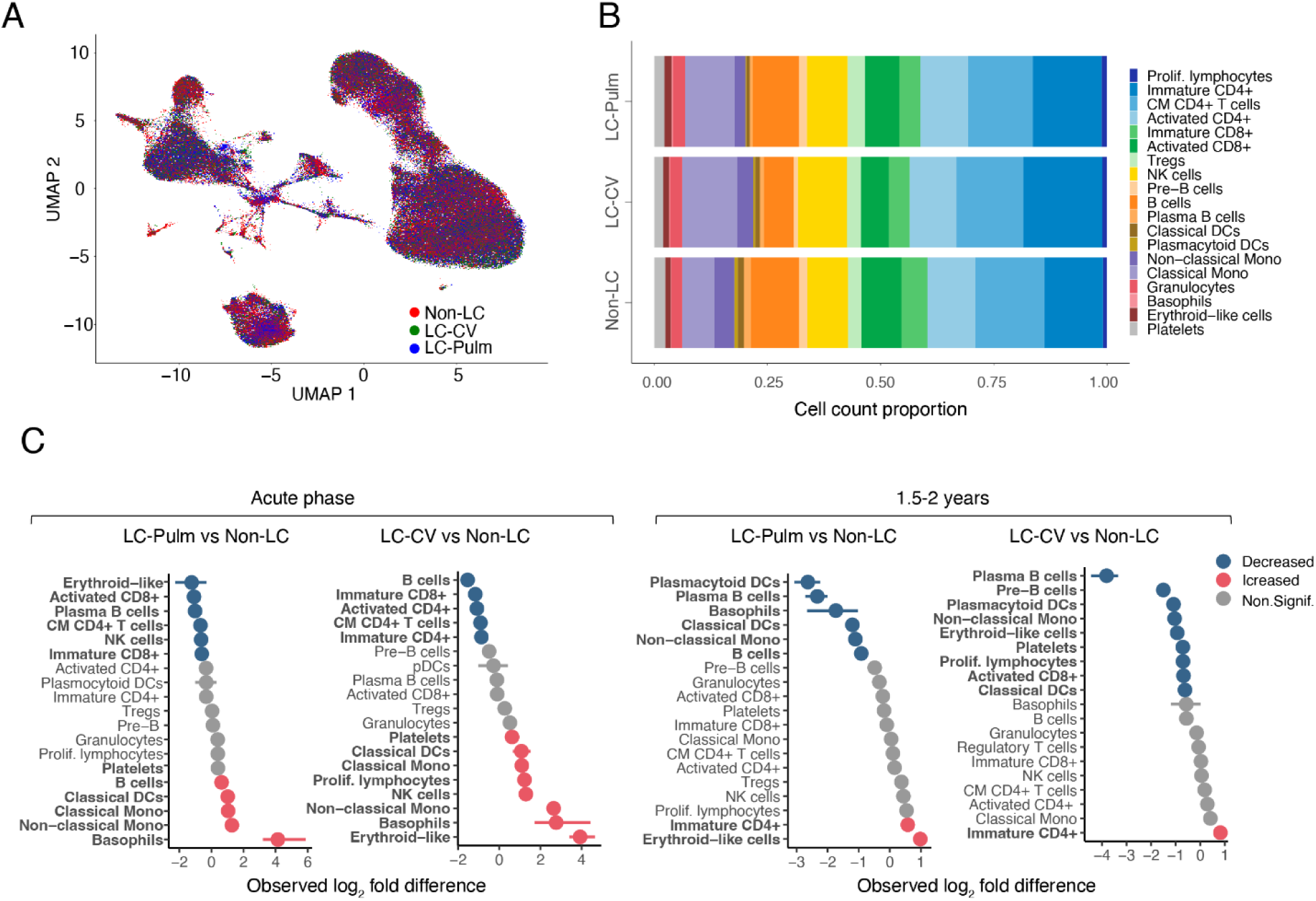
Proportional differences in identified cell populations across conditions and time points. A Overlay of the single cell landscape across three conditions: patients who did not develop Long-COVID (Non-LC, n = 3), patients who developed Long-COVID with cardiovascular (LC- CV, n = 3) and pulmonary (LC-Pulm, n = 3) complications, shown as a UMAP projection. Each point represents a cell, coloured according to their respective sample groups. B Distribution of cell population proportions in each of the three conditions. Sample complication groups are displayed along the y-axis, and cell count proportions are displayed along the x-axis. C Relative cell numbers in each cell population, compared between LC-Pulm and Non-LC, and between LC-CV and Non-LC patient groups at two time points: the acute phase of the COVID- 19 infection and 1.5 – 2 years later, expressed as log_2_ difference.

These clustering patterns can be further emphasized by observing the relative abundance of identified cell populations between conditions (Fig. 2B). The most pronounced differences were observed in Mono, T cell, and B cell repertoires. Non-classical Mono were the least abundant in the Non-LC group. In contrast, the classical Mono population was more prominent in the LC- Pulm group and even more so in the LC-CV group, suggesting a possible expansion of this population in patients who developed pulmonary or cardiovascular complications. Similarly, the population proportion of immature CD4^+^ T cells and proliferating lymphocytes was more prominent in both complication groups compared to the Non-LC group. Immature and activated CD8^+^ T cell populations were most abundant in the Non-LC group. Comparably, the plasma B cell population was more prominent in the fully recovered Non-LC group. Interestingly, both classical DCs and plasmacytoid DCs were more abundant in the Non-LC group, with the least abundance in the LC-Pulm group.

Comparison of the relative number of cells in the identified populations reveals evident changes at the acute phase of COVID-19 and even 1.5 – 2 years post-infection across disease conditions (Fig. 2C). During the acute phase, a significant increase in the relative number of basophils was observed in both complication groups, with the LC-Pulm group showing the highest observed log_2_ fold increase. Conversely, erythroid-like cell numbers increased most in the LC-CV group, whereas this population notably decreased in the LC-Pulm group at the same time point. Minor but considerable increases were observed in non-classical Mono, classical Mono and classical DCs in both study groups. B cell numbers were evidently higher in the LC- Pulm group, while the LC-CV patients showed a notable increase in Natural Killer (NK) cell and proliferating lymphocyte (prolif. lymphocyte) populations. A decrease in the relative number of immune cells during the acute phase was significant for multiple lymphoid cell populations in both Long-COVID complication groups. No notable differences in cell population proportions were observed for both study groups in plasmacytoid DCs, pre-B cells, T_regs_, granulocytes and platelets, as well as CD4^+^ and prolif. lymphocyte subsets in LC-Pulm, and plasma B and cytotoxic T cells in LC-CV (Fig. 2C, right panel).

At 1.5 – 2 years post-acute infection, only a limited number of cell populations showed a considerably increased relative number of cells in the complication groups. Immature CD4^+^ T cell numbers were elevated in both subject groups, while erythroid-like cell amount was increased exclusively in the LC-Pulm group. Long-COVID patients displayed a noticeable decline in plasma B cells and plasmacytoid DCs, with smaller reductions in B and pre-B cells, non-classical Mono, erythroid-like cells, platelets, prolif. lymphocytes, activated CD8^+^ T cells, classical DCs, and basophils. No distinct differences in cell numbers at 1.5 – 2 years post-infection were found among several populations, including most lymphocytes and classical Mono in LC-CV, and additionally platelets in LC-Pulm group.

To investigate cell-to-cell communication, we used CellChat analysis to infer the differential strength and number of interactions in cell populations of fully recovered (Non-LC) and Long-COVID (LC) patients. Initially, differential interaction analysis was performed for the absolute cell signalling in Non-LC and both LC study groups (Fig. 3A, B), and then within each cell population for the interaction strength (Fig. 3C) and number of interactions (Appendix Fig. S2) at the acute infection phase and 1.5 – 2 years post-infection. Significant differences were observed in cell-to-cell interaction patterns, with LC patient groups (LC-Pulm and LC-CV) exhibiting increased strength and number interactions compared to the Non-LC group at the acute phase of the COVID-19 infection (Fig. 3A). At 1.5 – 2 years post-infection, LC groups show a significant decrease in the overall cell-to-cell communication strength. In contrast, the number of interactions was significantly increased (Fig. 3B).

**Figure 3.**
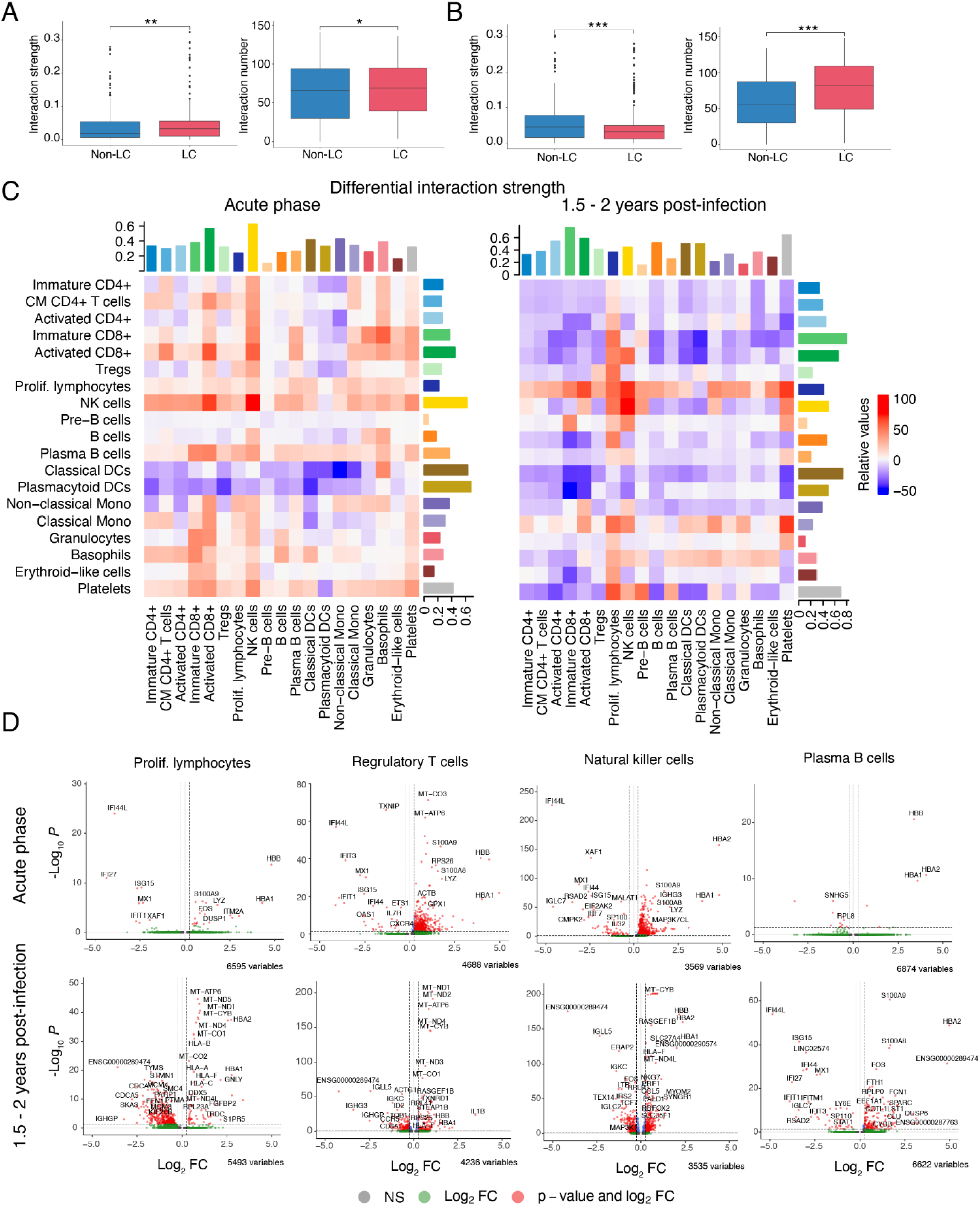
Intercellular communication between identified cell populations in recovered (Non-LC) and Long-COVID (LC) patient samples. A, B Total cell-to-cell interaction strength and number of interactions between patients who did (LC-CV and LC-Pulm groups) and did not develop Long-COVID (Non-LC group) at the time of acute infection (A) and 1.5 – 2 years later (B). C Heatmaps showing the differential interaction strength between previously identified cell populations. The y-axis represents the cell populations in which cells express the ligands of pathways from the CellChatDB Secreted Signalling database (sending), and cell populations where cells express the corresponding receptor (receiving) are depicted on the x-axis. Differential interaction strength is compared between fully recovered (Non-LC, n = 3) and Long-COVID (LC-Pulm and LC-CV, n = 6) patients during the acute phase of infection and 1.5 – 2 years post-infection (C). D Differential gene expression (x = Log_2_FC, y = -Log_10_P) between Non-LC and LC groups in prolif. lymphocyte, T_regs_, NK, and Plasma B cell populations at the time of acute infection and 1.5 – 2 years post-infection.

We further explored the observed decrease in interaction strength by individually analysing each of the previously identified cell populations, represented in Fig. 3C. High differential signal sending strength was observed in immature and activated CD8+ T cell, classical DCs and platelet populations, while lowest differential signalling capacity was found for pre-B cells and granulocytes at 1.5 – 2 years post-infection (Fig. 3C, 1.5 – 2 years post-infection). Likewise, the highest differential signal-receiving capacity was found for immature cytotoxic T cells and platelets, while the lowest signal-receiving strength was found for pre-B cell and granulocyte populations. Although the interaction strength was generally decreased across most cell populations at 1.5 – 2 years post-infection for LC patients compared with Non-LC, prolif. lymphocytes, classical Mono, NK cells, and basophils show increased interaction with multiple other cell populations, emphasizing the connection between proliferative lymphocyte and NK cell groups. Prolif. lymphocytes in the LC group consistently and robustly show an increase in interaction strength with all receiving cell populations, apart from plasmacytoid DCs, whereas classical Mono show increased interaction with all, except plasma-B cell populations.

On the other hand, during the acute phase of the infection, overall interaction strength is increased for LC groups when compared with the Non-LC group (Fig. 3C, acute phase). The highest increase in outgoing signalling strength was observed for NKs, the highest decrease - classical and plasmacytoid DCs, while the signal-receiving capacity was highest for activated CD8+ T and NK cell populations. Pre-B cells show the lowest change in receiving and sending signal strength, which remains low at the post-infection stage as observed previously. The strength of interactions at the acute phase was reduced for classical DCs and plasmacytoid DCs with all receiving cell populations except basophils. Moreover, there was an apparent decrease in interaction strength capacity between T cell subsets with DCs and non-classical Mono.

Lastly, we looked at DEGs in prolif. lymphocyte, T_reg_, NK, and Plasma B cell populations between Non-LC and LC groups (Fig. 5D). Both the acute phase and 1.5 - 2 years post-infection haemoglobin-associated gene expression is significantly upregulated (*HBB, HBA1, HBA2*) in all cell populations, suggestive of a sustained haemoglobin production dysfunction, which has been observed in COVID-19 patients previously (Zhang *et al*, 2022). Likewise, all cell populations at both time points, apart from NKs at 1.5 - 2 years post-infection, show upregulation of *S100* family genes (*S100A8, S100A9*), which have been implicated in playing a role in Long-COVID (Boucher *et al*, 2024; Deguchi *et al*, 2021; Mellett & Khader, 2022). Furthermore, all cell populations of Long-COVID patients in the acute phase show significantly downregulated IFN-I associated genes (*IFI44L, IFI44 IFI27, IFIT1, IFIT3*) and *ISG15* (IFN-stimulated gene 15), essential to viral infection defence (Munnur *et al*, 2021; Sarkar *et al*, 2023), which is highly suggestive of an impaired IFN-I response. Interestingly, T_regs_ show upregulation of *IL1B* (Interleukin 1-β) 1.5 - 2 years post-infection. This cytokine has been found to be elevated in severe COVID-19, and the expression at a later stage may indicate ongoing pro-inflammatory cytokine involvement (Mardi *et al*, 2021). Remarkably, prolif. lymphocytes in LC groups at 1.5 to 2 years post-infection highly expressed a range of Human Leukocyte Antigen (*HLA*) genes (*HLA-A, HLA-B, HLA-C, HLA-F*), which is particularly intriguing, as these findings may offer clues into the determinants influencing the emergence of symptoms associated with Long-COVID (Xie *et al*, 2024; Hoseinnezhad *et al*, 2024; Augusto *et al*, 2023). Together, these findings suggest that immune system dysfunction begins during the acute phase and continues up to 1.5 – 2 years after infection in patients with Long-COVID.

Next, we conducted an in-depth analysis of specific immune cell subsets to explore further the underlying mechanisms driving differential cell signalling patterns. We assessed the potential COVID-19-induced immune regulatory effects by examining the properties of T, NK, and proliferative lymphocyte subsets. We compared the expression of well-known cytotoxicity (*IFNG, GNLY, GZMA, GZMH, KLRK1, KLRB1, CTSW, CST7*) (Fig. 4A) and exhaustion-associated (*TIGIT, PDCD1, HAVCR2, TOX, IRF4, LAG3, BTLA, VSIR, CD96, CD28, CD226*) (Fig. 4B) genes to describe cell state variance between fully recovered (Non-LC) and LC study groups (LC-CV and LC-Pulm).

**Figure 4.**
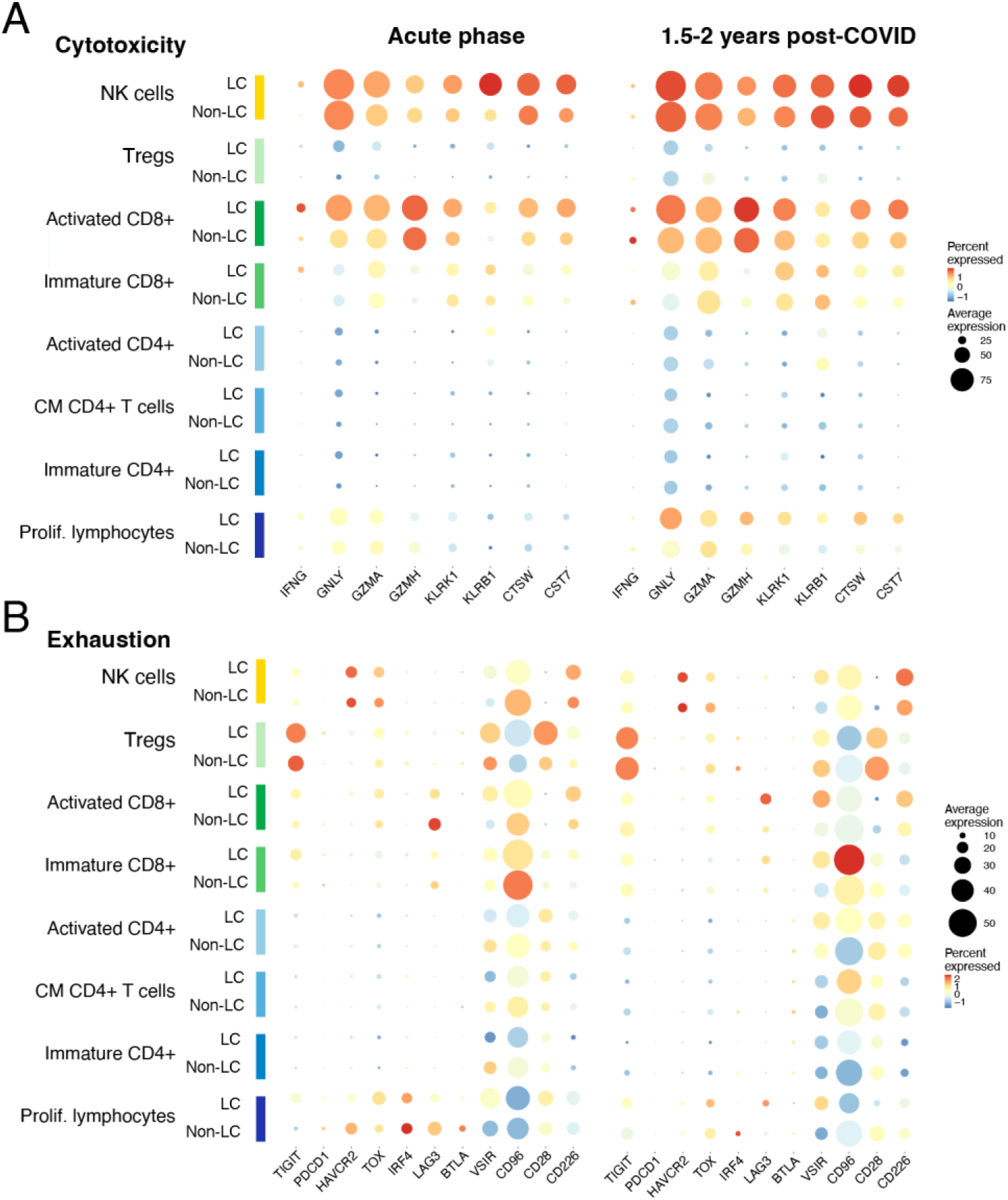
Cytotoxicity and exhaustion cell states. A, B Well-known (A) cytotoxicity (*IFNG, GNLY, GZMA, GZMH, KLRK1, KLRB1, CTSW, CST7*) and (B) exhaustion-associated (*TIGIT, PDCD1, HAVCR2, TOX, IRF4, LAG3, BTLA, VSIR, CD96, CD28, CD226*) gene expression in T, NK and proliferative lymphocyte subsets during the acute phase of infection and 1.5 – 2 years post-infection for fully-recovered (Non-LC) and complication (LC) patient groups. Coloured cell populations and patient groups are represented on the y-axis, with genes of interest shown along the x-axis. Average gene expression is indicated as the dot size, and the dot colour denotes the percentage of cells in which the gene of interest was expressed.

During the acute phase of infection, NK cells highly expressed cytotoxicity markers, particularly *KLRB1* (Killer Cell Lectin-Like Receptor B1, also called *CD161*) (Kurioka *et al*, 2018) in LC groups. Additionally, *CST7*, which encodes Cystatin-F and is known to modulate the cytotoxic activity (Perišić Nanut *et al*, 2017), expression was higher in both NK and activated CD8^+^ T cells in LC groups during the acute phase. Activated CD8^+^ T cells showed high average expression of a gene encoding Cathepsin W (*CSTW*) and *CST7* in LC groups at the acute phase, remaining highly expressed 1.5 – 2 years post-infection. Importantly, proliferative lymphocytes also displayed sustained upregulation of cytotoxicity markers *GNLY, GZMH* (Granzyme H)*, KLRK1* (Killer Cell Lectin-Like Receptor K1)*, KLRB1, CST7* and *CSTW* 1.5 – 2 years post-infection (Fig. 4A). The observed increased cytotoxic activity 1.5 – 2 years post-infection indicates that these cells undergo prolonged activation and possible subsequential exhaustion.

Exhaustion-associated patterns can also be observed in LC groups across the time points. T_regs_ at the acute phase express *TIGIT* (T Cell Immunoreceptor With Ig And ITIM Domains) and *VSIR* (V-Set Immunoregulatory Receptor), a T cell inhibitory receptor (Lines *et al*, 2014) which encodes V-domain Ig suppressor of T cell activation protein (VISTA). As well, T_regs_ in the LC group showed higher expression of *CD28*, a co-stimulatory receptor (Hui *et al*, 2017; Strioga *et al*, 2011), during the acute phase. An activation-associated marker, *CD96* or *Tactile* (T cell activation, increased late expression) (Wu *et al*, 2022), was highly expressed in immature CD8^+^ cells in the Non-LC group during the acute phase. The expression of this marker was even more pronounced in the LC group at 1.5 – 2 years post-infection. Activated CD8^+^ cells expressed *LAG3* (Lymphocyte Activating Gene 3) in the Non-LC group, but this effect seems reversed over time, as the LC group exhibits higher *LAG3* expression at 1.5 – 2 years post-infection. At 1.5 – 2 years post-infection, proliferative lymphocytes, activated CD8^+^ T cells, and NK cells in the LC group continued to express *VSIR* at higher levels than the respective populations in the Non-LC group (Fig. 4B). This reinforces the notion that immune regulation is uniquely altered in COVID- 19 patients, which may persist for long after the initial infection resolution.

To better acknowledge the possible physiological implications of the previously observed cellular variations between the conditions, we calculated cell scores for IFN-α, β, and γ response, migration, and apoptosis-associated gene expression in T, NK, B cell and Mono subpopulations (Fig. 5.A), and across Non-LC and LC groups as a whole (Fig. 5B) at the acute and post-infection time points.

**Figure 5.**
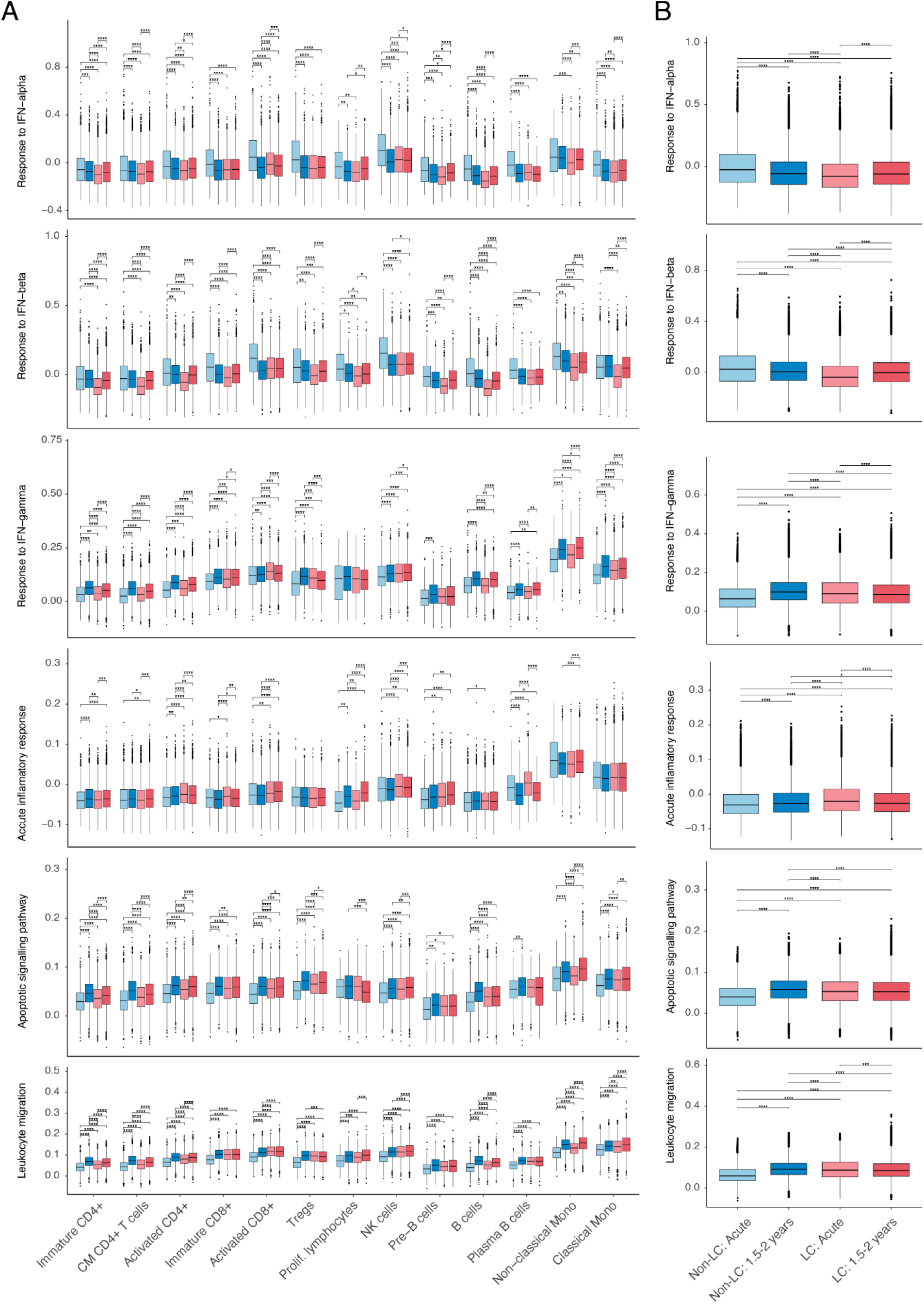
T, NK, B cell, and Mono subpopulation cell state scores between conditions and time points. A, B Cell state scores calculated using AddModuleScore function in Seurat indicating the IFN-α, IFN-β, IFN-γ response, inflammatory response, apoptosis, and migration-associated gene expression in (A) each of the T, NK, B cell and Mono subpopulations, and (B) total scores between Non-LC and LC groups at the acute phase of the infection, as well as at 1.5 – 2 years post-infection.

During the acute phase, IFN-α and β responses were significantly higher in the Non-LC group across all analysed subpopulations, whereas IFN-γ response was elevated in LC cells, particularly in T cells, NK cells, and Mono (Fig. 5A), together indicating a stronger type II, but weaker type I interferon (IFN-I) response in LC patients (Fig. 5B). Acute inflammatory response and apoptotic signalling associated gene expression were higher in several lymphocyte populations, while leukocyte migration was higher among all analysed cell subpopulations in the LC group (Fig. 5A, B). Taken together, these findings point to a heightened inflammatory immune response in patients who go on to develop Long-COVID during the acute infection, suggesting early onset immune system perturbations.

At 1.5 – 2 years post-infection, the IFN-α response was higher in certain LC lymphocyte subpopulations, including immature CD4^+^, activated CD8^+^, proliferating lymphocytes, NK cells, and pre-B cells. However, IFN-α response was lower in non-classical Mono, and no significant differences were observed in total IFN-α response between the study groups. In contrast, IFN-β response remained lower in LC for most subpopulations, except in activated CD8^+^ T cells, where it was higher. IFN-γ response was generally reduced in LC, except for certain populations, such as activated CD8^+^ T cells, NK cells, and non-classical Mono, where it remained elevated. Inflammatory response was persistently higher in the LC group across multiple lymphoid cell types even 1.5 – 2 years post-infection. At the same time, apoptotic signalling was lower in most T and B cells, and proliferating lymphocyte populations in LC, but remained significantly higher in NK cells and non-classical Mono. Meanwhile, leukocyte migration was elevated in immature and activated CD4^+^ and B cells in LC but reduced in activated CD8+, NK, and Mono subpopulations (Fig. 5A). In total, acute inflammatory response and apoptotic signalling were significantly lower, but leukocyte migration, IFN-β and γ responses were higher in LC patients (Fig. 5B), indicating an ongoing immune system activation 1.5 – 2 years post-infection.

## Discussion

Here, we present a unique dataset with a detailed single-cell transcriptomic analysis of PBMCs in both acute and Long-COVID patients. Our findings illustrate shifts in the immune cell subpopulations between groups, cell communication alterations, and differential expression of key function genes. These collective findings provide insights into the immune landscape of COVID-19 as the disease progresses into a chronic, Long-COVID state. Several cell groups stand out in the all-encompassing analyses: prolif. lymphocytes, T cells, NKs, Mono, and B cells. These patterns indicate that distinct blood immune cell populations may play specific roles in the progression and persistence of Long-COVID.

Our analysis reveals LC patient cells express increased exhaustion-associated markers, like *LAG3* and *VSIR*, suggesting long-lasting immune regulation changes in Long-COVID patients. Whether this plays a role in the progression or persistence of Long-COVID remains to be studied, however, the presented findings provide compelling evidence to support this possibility. Together, these exhaustion-associated gene expression profiles across T cell subsets and NK cells suggest a potential immune system anergy in Long-COVID, highlighting an area for further exploration. Moreover, prolif. lymphocytes and multiple T cell subsets exhibit increased cytotoxic activity, which may exacerbate tissue damage and drive chronic inflammation. The expression *HLA* genes 1.5 – 2 years post-infection further proposes prolif. lymphocyte involvement in Long-COVID persistence and pathogenesis.

The apparent increased signalling patterns in prolif. lymphocytes, along with the upregulation of cytotoxicity-associated genes observed during the Long-COVID stage, is suggestive of persistent immune activation, potentially linked to incomplete recovery from the acute phase of infection. This is particularly compelling because even though no changes were observed in the acute stage, 1.5 – 2 years post-infection, IFN-α and acute inflammatory responses were higher in prolif. lymphocytes, along with increased intercellular signalling strength in Long-COVID patients. These observations outline prolif. lymphocytes as one of the potential key modulators of Long-COVID complications.

Both Long-COVID complication groups show relatively larger Mono populations at the acute phase, which, coupled with high IFN-γ response, may indicate a stronger necessity for functional cell recruitment in patients who develop Long-COVID. Conversely, a decrease in the non-classical Mono population 1.5 – 2 years post-infection, along with low IFN-α, IFN-β, and IFN-γ responses in the Long-COVID patient group, signifies the involvement of the innate immune system regulation. Mononuclear phagocyte role has been implicated in contributing to severe COVID-19 cases (Edahiro *et al*, 2023) and Long-COVID pathogenesis (Cervia-Hasler *et al*, 2025)

B cells to show a reduced ability to interact with multiple cell populations in Long-COVID patient groups, which raises concern about long-term adaptive immunity function. While at the acute phase, pre- and plasma B cells exhibit more prominent inflammatory responses, at 1.5 – 2 years post-infection B cell inflammatory properties (IFN-β and IFN-γ response) were lower in Long-COVID patients versus Non-LC. The diminished ability of pre-B cells to interact with other immune cell populations, their reduced inflammatory properties 1.5 – 2 years post-infection may affect immune memory and warrant further investigation.

While our single-cell RNA analyses provide a detailed overview of the population dynamics and molecular processes within blood immune cells, functional assays, flow cytometric analysis and proteomics, can further exemplify the observed differences and whether these can be targeted for Long-COVID treatment. Our study has revealed interesting observations about the prolif. lymphocyte subpopulation, a potential area for independent exploration with more targeted approaches. The sample size, though comprehensive, may not fully reflect the Long-COVID manifestations in patients with a more diverse range of symptoms, including neurological complications. However, our dataset is unique in its focus on a well-characterized cohort of initially hospitalized patients with prolonged Long-COVID, extending beyond the typical study periods and offering a rare insight into the condition’s long-term effects (Peter *et al*, 2025; Sivan *et al*, 2025). It would be valuable to explore how the changes observed in PBMCs reflect broader systemic immune responses. To address this, we aim to integrate the results of this study with previously published gut microbiome data from patients within the same cohort (Brīvība *et al*, 2024) which will facilitate the ongoing research into the pathophysiology of Long-COVID, helping to unravel its underlying mechanisms and systemic consequences.

It is essential to note that Long-COVID manifests with a variety of symptoms at different levels of severity, and issues with classification and standardization for this diagnosis may mean cases remain overlooked. The interplay between Long-COVID symptoms and immune recovery remains poorly understood, with the determinants and triggers underlying the development of Long-COVID still under investigation (Peter *et al*, 2025). This raises the possibility that while immune dysregulation is evident, it may not always translate into clinical diagnosis.

Taken together, this study shows distinct clustering of cell types, mitigated intercellular communication, and altered cell states in Long-COVID patients. Immune cell dysregulation in Long-COVID is evident in our findings and previous studies (Yin *et al*, 2024; Cervia-Hasler *et al*, 2025; Scott *et al*, 2023). Differentially expressed genes and the involvement of the innate immune system regulation, though widely documented in severe acute COVID-19 cases, may play a part in the development or persistence of Long-COVID, as highlighted by the outcomes of this investigation. The detailed comparison of cell states, populations, and ligand-receptor signalling provides a compelling outlook on the immune processes and cell states in acute and Long-COVID.

## Materials and Methods

### Methods and Protocols

#### Overview of the patient cohort

The prospective study involved female patients who had been hospitalized during acute COVID- 19 (average age: 55, median: 52; average BMI: 28,22, median: 27,72) at Riga East University Hospital, Latvia between October 2020 to January 2021. Written informed consent was obtained from every participant before their inclusion in the study, the study was approved by the Central Medical Ethics Committee of Latvia (No. 01-29.1.2/928) and carried out according to the Declaration of Helsinki and the Department of Health and Human Services Belmont Report. COVID-19 infection was confirmed by RT-PCR test, and blood sampling was arranged at the acute infection phase, at 3 months, and between 18 and 24 months post-acute infection. Peripheral blood samples were obtained using BD Vacutainer® CPT™. 18-24 months post-acute COVID-19, patients who displayed Long-COVID with cardiovascular (LC-CV, n = 3, average age: 54, median: 52; average BMI: 34,56, median: 33,09) and pulmonary complications (LC-Pulm, n = 3, average age: 65, median: 68; average BMI: 24,56, median: 23,88) along with patients who did not develop Long-COVID (Non-LC, n = 3; average age: 47, median: 47; average BMI: 25,57, median: 27,72), were selected for this study. All patients are non-smokers, and two patients reported using antiviral medicine (Remdesivir), as well as most used anticoagulant medicines (Enoxaparin) during the acute phase of COVID-19. All patients except one in each complication group (LC-CV and LC-Pulm) have been vaccinated with Pfizer-BioNTech, Moderna or Jannsen vaccines (Appendix Table S1).

#### Sample preparation and storage

Peripheral blood samples were collected using the BD Vacutainer® CPT™ collection tubes according to the manufacturer’s instructions and processed less than two hours after sample collection to preserve cell quality. All safety measures when working with human samples and pathogens were taken into consideration out throughout the study. Mononuclear cells were then isolated using the density gradient centrifugation method, counted using Trypan blue (Thermo Fisher Scientific), and stored in Foetal Bovine Serum (FBS) (Thermo Fisher Scientific) with added 10% dimethyl sulfoxide (DMSO) (Sigma-Aldrich) in liquid nitrogen.

#### Single-cell RNA library preparation

Single-cell suspensions were washed twice with Phosphate Buffered Saline (PBS) with 0.04% Bovine Serum Albumin (BSA) (Sigma-Aldrich), cells were counted, and 30 000 cell input was used for further analysis with MGI DNBelab C Series High-throughput Single-cell RNA Library Preparation Set V2.0 (MGI, Shenzhen, China). In short, single cells were combined with magnetic beads and encapsulated in oil droplets using the DNBelab C4 microfluidic device for mRNA hybridization. Magnetic beads were collected, mRNA from single cells was reverse transcribed, and libraries were prepared according to the manufacturer’s protocol.

The resulting cDNA product fragment size was distributed between 1149 bp to 1492 bp. cDNA and oligo libraries had fragment sizes between 428 bp to 515 bp and 181 bp to 190 bp, respectively. Quality control of fragment size and concentration was performed using Agilent 2100 Bioanalyzer System and Qubit dsDNA HS Assay kit for Qubit® 2.0 Fluorometer (Thermo Fisher Scientific).

#### Sequencing

Sequencing was carried out using DNBSEQ-G400RS platform (MGI, Shenzhen, China). Prepared libraries were circularized, and DNA NanoBalls (DNBs) were generated according to the manufacturer’s protocol. The resulting DNBs were loaded onto the sequencing flow cell and sequenced with DNBSEQ-G400RS High-throughput Sequencing kit FCL PE100 (MGI, Shenzhen, China) according to the manufacturer’s instructions.

#### Data analysis

Raw sequencing data processing, which includes primary quality control, alignment, annotation, and quantification, was done using the DNBelab C Series HT scRNA analysis software (*v2.1.1*) (MGI Tech bioinformatics R&D). Within this pipeline, the sequence reads were aligned to the GENCODE v44 (Mudge *et al*, 2025) Homo sapiens reference genome. The filtered expression matrices and cell barcodes were then used for the downstream analysis in the R (R Core Team, 2021) /RStudio (*v4.4.2*/*v2023.12.1*) (RStudio Team, 2020) environment using the Seurat (*v5.1.0*) (Hao *et al*, 2024) workflow.

The downstream analysis consisted of two parts. After importing the data into the Seurat ecosystem, we started with 39 349 genes across 223 087 cells. After running all the steps described below until the cell type detection, we determined that we had 44 369 cells in three clusters for whom we could not determine their specific cell populations due to their low overall expression levels and no specific marker signatures for the known immune cell types. Therefore, they were removed from the data set, and all the initial data steps were repeated. After eliminating these cells, we continued with the remaining 178 719 cells.

We first started by examining the distributions for multiple cell and gene quality indicators. Based on these results, we removed all cells with less than 500 genes, mitochondrial gene ratio > 20%, novelty score (log_10_GenesPerUMI) < 85%, and which had less than 500 or more than 5000 unique molecular identifiers. Afterwards, we also removed genes that were expressed in less than 10 cells. The additional quality filtering resulted in 38 280 genes across 174 336 cells.

Next, we scaled our data and selected the 3000 most variable features using the “vst” selection method. Subsequently, we calculated the cell cycle scores and evaluated the cell grouping across multiple variables using the principal component analysis (PCA) method to select the regression variables for normalization. SCTransform was used for the sample level normalization with the mitochondrial ratio, cell cycle S, and G2M scores used as the regression variables.

To integrate the data, we selected 3000 integration features on which we then performed PCA ordination and subsequent integration on the SCT data using the Harmony (*v1.2.0*) (Korsunsky *et al*, 2019) method where the reduction method was specified as PCA with the Long-COVID-19 complication groups and time points provided as covariates. Based on the elbow plot generated on the cumulative explained variation of the principal components (PCs), we selected the first 30 PCs for the uniform manifold approximation and projection (UMAP) dimension reduction and to also determine the clusters at different clustering resolutions (0.2 - 2.0) with the Shared Nearest Neighbor (SNN) method, from which we chose the resolution of 0.8, as it provided a reasonable number of clusters together with good visual separation when evaluating the UMAP plot. The resulting cluster resolution yielded 22 cell clusters.

#### Cell type annotation

To determine the cell types in our dataset, we first started with the ScType (*v1.0*) (Ianevski *et al*, 2022) approach, which uses a precompiled list of cell type markers for multiple cell types. Using this approach, we obtained a preliminary list of cell populations present in our dataset. Next, we evaluated the ScType results and used the Seurat’s FindAllMarkers method with the Wilcoxon test to obtain a list of differentially expressed markers (min.pct = 0.25, l2fc > 0.25) between the determined cell clusters along with our own list of cell markers. The differentially expressed markers were examined using a DotPlot graph for their prevalence in the proportion of cells in each cluster and their expression levels. As a final result, we obtained a list of 19 unique cell populations for the 22 clusters detected.

To determine the differences in cell counts between different time points and complication groups for the detected cell types, we used a cell proportion test from the scProportionTest (*v0.0.0.9000*) (Miller *et al*, 2021) package and proportionally transformed cell count bar plot graphs for each cell type.

Cell exhaustion and cytotoxicity states were defined for proliferating lymphocytes, T and Natural Killer (NK) cell subpopulations using well-known marker genes associated with either cell exhaustion (*TIGIT, PDCD1, HAVCR2, TOX, IRF4, LAG3, BTLA, VSIR, CD96, CD28, CD226*) or cytotoxicity (*IFNG, GNLY, GZMA, GZMH, KLRK1, KLRB1, CTSW, CST7*). AddModuleScore function was used in Seurat to establish a score for each T, NK, B cell and monocyte subsets indicating the interferon (IFN)-α/β/γ response, inflammatory response, apoptosis, migration scores, using RESPONSE TO INTERFERON ALPHA (GO:0035455), RESPONSE TO INTERFERON BETA (GO:0035456), ACUTE INFLAMMATORY RESPONSE (GO:0002526), APOPTOTIC SIGNALING PATHWAY (GO:0097190) and LEUKOCYTE MIGRATION (GO:0050900) gene sets, respectively. The exhaustion scores were compared using the Kruskal-Wallis omnibus test to evaluate the presence of overall differences and the pairwise Dunn test with Holm-Bonferroni p-value adjustment from the rstatix (*v0.7.2*) (Kassambara, 2023a) package to obtain results for individual pairs of comparisons. The results were visualized in the form of box plots with the help of ggplot2 (*v3.5.1*) (Wickham *et al*, 2016) and ggpubr (*v0.6.0*) (Kassambara, 2023b) packages.

#### Differential gene expression

Differential expression (DE) analysis between cell types of interest in the Non-LC and Long-COVID cohorts, as well as Non-LC vs. pulmonary (LC-Pulm) and cardiovascular (LC-CV) complication groups separately, was performed using Seurat’s FindAllMarkers function with the Wilcoxon test option and the cut-off values of p-adj < 0.05 and log_2_ Fold Change (FC) > = 0.25. Only genes expressed in at least 10% of cells in the cell group were considered. Upon completion of DE analysis, the results were visualized in volcano plots using the EnhancedVolcano (*v1.24.0*) package (Kevin *et al*, 2024).

#### Differential intercellular communication inference with CellChat

To provide a more systems-level approach to studying the cellular dysregulation associated with Long-COVID, the CellChat (*v2.1.1*) (Jin *et al*, 2021) program was used to infer cell to cell communication patterns and the changes thereof between the complication and control groups from scRNA-seq gene expression data. CellChat employs an extensive database of ligand-receptor interactions (CellChatDB) combined with mass-action-based models for statistical inference of communication probabilities between cell groups that adjusts for cell population size. With this communication probability inference, CellChat represents the scRNA-seq dataset as a network of interacting cell groups, which can then be projected onto the human protein-to-protein interaction network for greater accuracy of results.

In this study the Secreted Signalling database from CellChatDB, which focuses on cell-to-cell communication through cytokine and chemokine signalling, was used. The integrated Seurat object of Long-COVID and Non-LC patient cells was converted into separate CellChat objects for each time point and condition. As per CellChat guidelines, overexpressed genes were identified at the threshold of p < 0.05, and expression data were projected onto the human protein-to-protein interaction network. Cell communication probabilities were calculated with the triMean type of average per-cell group expression estimation and filtered at the threshold of at least 10 cells in each communicating group. Based on the calculated probabilities, cell-to-cell communication networks were then constructed and aggregated. Long-COVID and Non-LC CellChat objects were merged for each visit and cellular communication analyses were performed following CellChat guidelines. Differential interaction number and strength per pair of cell types between the patient groups were visualized as a heatmap (Fig. 3C., Appendix Fig. S2) using the netVisual_heatmap function. The changes in total mean interaction number and interaction strength were also compared between Long-COVID and Non-LC patient groups at the acute phase and 1.5 – 2 years post-infection using a two-tailed Wilcoxon rank-sum test (p < 0.05 significance level).

## Acknowledgements

The authors acknowledge the Latvian Biomedical Research and Study Centre and the Genome Database of the Latvian Population for providing the infrastructure, biological material and data. The authors also acknowledge SIA Latvia MGI Tech for providing technical support throughout the study.

## Author contributions

**Marta Līva Spriņģe:** manuscript writing; investigation; conceptualisation; revision; data interpretation; methodology**. Kristīne Vaivode:** visualisation; conceptualisation; revision; data interpretation; methodology; supervision. **Rihards Saksis:** visualisation; data analysis; data interpretation; writing and revision. **Nineļa Miriama Vainšeļbauma:** visualisation; data analysis; data interpretation; writing and revision. **Laura Ansone:** patient recruitment; data acquisition. **Monta Brīvība:** project administration; data acquisition. **Helvijs Niedra:** methodology; study design; supervision. **Vita Rovīte:** conceptualisation; methodology; study design; project administration; funding acquisition; resource allocation; supervision.

## Funding

This research was funded by the European Regional Development Fund (ERDF), Measure 1.1.1.1 “Support for Applied Research”, grant number 1.1.1.1/21/A/029. Rihards Saksis was supported by the project “Strengthening of the Capacity of Doctoral Studies at the University of Latvia within the Framework of the New Doctoral Model”, identification No. 8.2.2.0/20/I/006. Laura Ansone and Helvijs Niedra (ANM_K_DG_27) was supported within the framework of the European Union’s Recovery and Resilience Mechanism project No.5.2.1.1.i.0/2/24/I/CFLA/001 “Consolidation of the Latvian Institute of Organic Synthesis and the Latvian Biomedical Research and Study Centre”.

## Disclosure and competing interest statement

The authors declare no conflict of interest.

## The Paper Explained

### Problem

Long-COVID aetiology remains poorly understood, but emerging evidence points to immune dysregulation as a potential mechanism involved in its development or persistence. As the number of individuals requiring long-term post-COVID-19 care continues to rise, understanding the immune responses over time is crucial. The current lack of prognostic tools and effective treatment options for Long-COVID underscores the necessity of a detailed investigation in patients with this debilitating chronic condition.

### Results

Our study provides a detailed investigation of the immune landscape, following patients from the acute phase into 1.5 – 2 years post-COVID-19. Using a scRNA-seq approach, we are able to characterize peripheral blood mononuclear cells (PBMCs) from patients with pulmonary and cardiovascular Long-COVID complications. By comparing immune cell populations and gene expression profiles between acute and post-acute phases, we identify key cell subpopulations, such as proliferating lymphocytes, monocytes, NK and T cell subsets, in which we observe immune dysregulation patterns. Our results show early changes in the immune landscape at the acute phase of infection in individuals who will go on to develop Long-COVID, including differential interferon responses and cell-to-cell communication. As well, we outline exhaustion and cytotoxicity associated gene expression, altered inflammatory signalling and intercellular communication as prominent patterns in Long-COVID at 1.5 – 2 years post-infection.

### Impact

Our work provides a rare insight into the lasting immune dysregulation 1.5 – 2 years post-acute COVID-19 infection, highlighting specific cell subpopulations involved and outlining potential modulators driving persistent symptoms. The study contributes to the ongoing efforts in elucidating the underlying mechanisms of Long-COVID pathogenesis and further underscores the importance of long-term monitoring of Long-COVID patients, as well as the necessity for further exploration and research into prognostic, diagnostic and therapeutic targets for Long-COVID management.

### Data Availability Section

ScRNA-seq data have been submitted at the Sequence Read Archive (PRJNA1208100) and will be deposited at Gene Expression Omnibus.

## Appendix

**Appendix Table S1.** Patient cohort characteristics.

| Patient | Study group | Age | Body mass index (BMI) | Vaccination (in order) | Medication during the acute phase of infection | How many times had COVID-19 |
| --- | --- | --- | --- | --- | --- | --- |
| P711 | LC-Pulm | 45 | 26,83 | Pfizer-BioNTech<br>Pfizer-BioNTech | Paracetamol<br>Diclofenac<br>Codeine<br>Ceftriaxone<br>Doxycycline<br>Chloropyramine hydrochloride<br>*Sirupus Pini Compositus*<br>Dexamethasone<br>Enoxaparin<br>Ringer's solution | 2 |
| P921 | LC-Pulm | 52 | 43,75 | Pfizer-BioNTech<br>Pfizer-BioNTech<br>Pfizer-BioNTech<br>Pfizer-BioNTech | N/A |  |
| P908 | LC-Pulm | 66 | 33,09 | N/A | N/A | N/A |
| P713 | Non-LC | 47 | 27,72 | Pfizer-BioNTech<br>Pfizer-BioNTech<br>Pfizer-BioNTech | Paracetamol<br>Codeine<br>Doxycycline<br>Bromhexine<br>Enoxaparin<br>Ascorbic Acid 10%<br>Omeprazole | 1 |
| P865 | Non-LC | 46 | 28,52 | Pfizer-BioNTech<br>Pfizer-BioNTech | Paracetamol<br>Ceftriaxone<br>Doxycycline<br>Remdesivir<br>Bromhexine<br>Enoxaparin<br>Ascorbic Acid 10%<br>Metoclopramide<br>Pancreatin<br>Neurorubine | 1 |
| P407 | Non-LC | 49 | 20,45 | Pfizer-BioNTech<br>Moderna | N/A | 1 |
| P841 | LC-CV | 55 | 28,34 | YES | Oxygen supplementation<br>Analgin<br>Ceftriaxone<br>Doxycycline<br>Remdesivir<br>Bromhexine<br>Dexamethasone<br>Enoxaparin<br>Ascorbic Acid 10%<br>Syr. Lactulose | 3 |
| P843 | LC-CV | 68 | 21,45 | NO | Oxygen supplementation<br>Analgin<br>Paracetamol<br>Ceftriaxone<br>Doxycycline 1<br>Bromhexine<br>Dexamethasone 8<br>Enoxaparin<br>Ascorbic Acid 10%<br>Omeprazole<br>Bromazepam | 2 |
| P854 | LC-CV | 71 | 23,88 | Janssen | Oxygen supplementation<br>Ceftriaxone<br>Doxycycline<br>Bromhexine<br>Dexamethasone<br>Enoxaparin<br>Ascorbic Acid 10%<br>*Mixturae nervinae*<br>*Sirupus Pini Compositus*<br>Quetiapine<br>Omeprazole | 1 |

**Appendix Figure S1.**
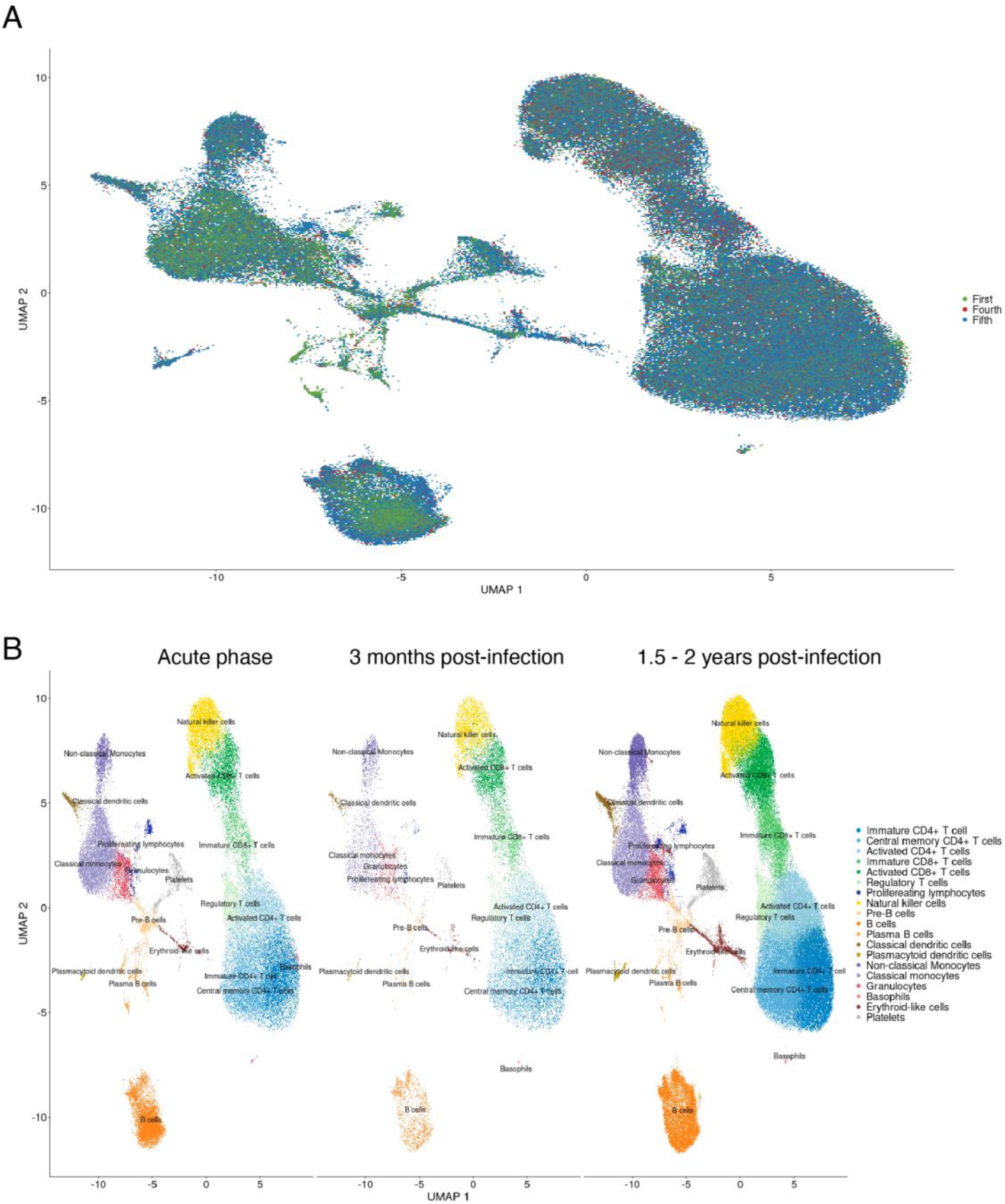
Immune cell landscape at acute COVID-19 infection, 3 months and 1.5 – 2 years post-infection. A UMAP overlay of all patient cells at the three timepoints: acute COVID-19 infection, 3 months and 1.5 – 2 years post-infection. B UMAP of the all patient cells at the three timepoints: acute COVID-19 infection, 3 months and 1.5 – 2 years post-infection.

**Appendix Figure S2.**
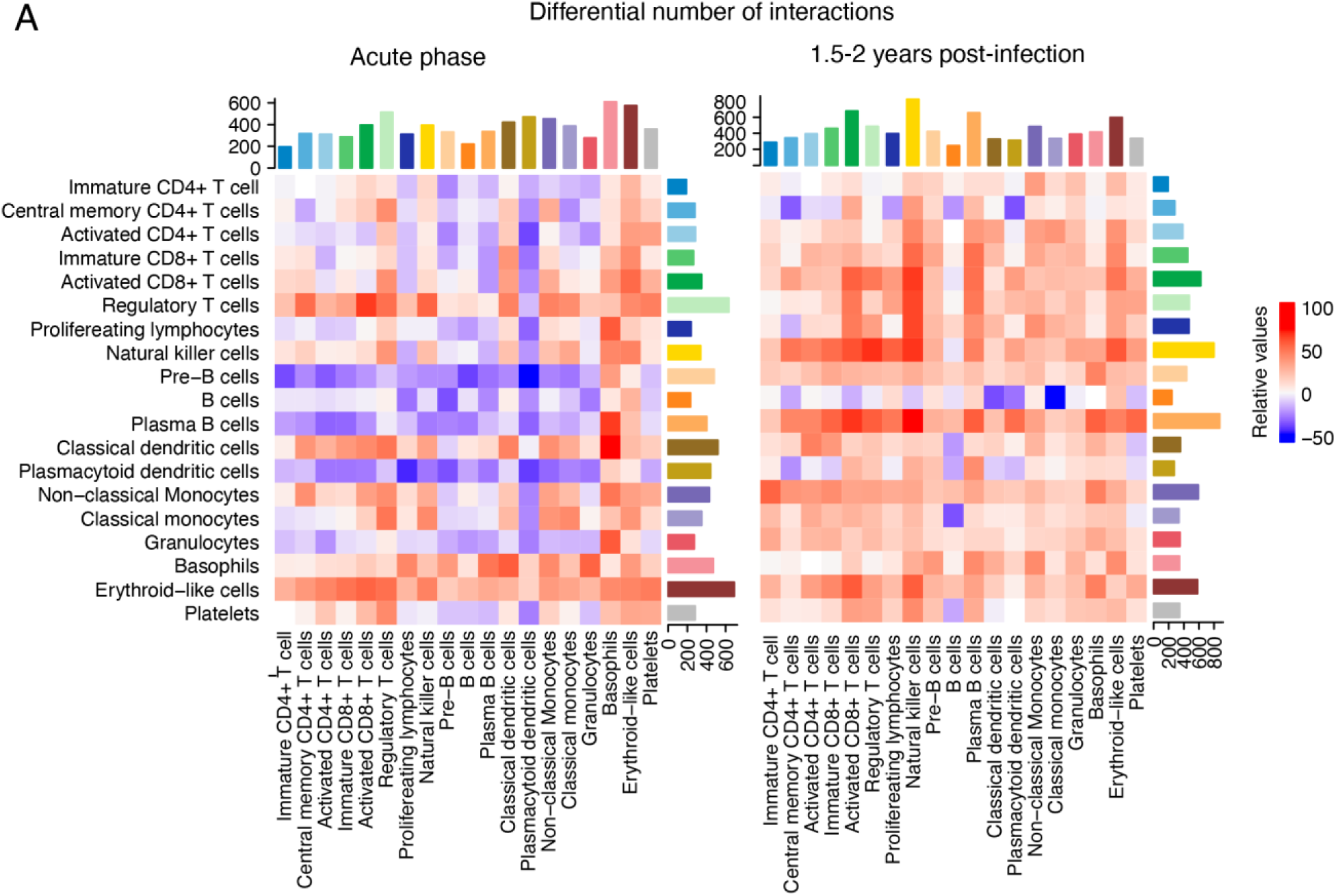
Cell-to-cell differential number of interactions between conditions at the acute phase and 1.5 – 2 years post-infection. A Heatmaps showing the differential number of interactions and in the identified cell populations, compared between fully recovered (Non-LC, n = 3) and Long-COVID (LC-Pulm and LC-CV, n = 6) patients during the acute phase of infection and 1.5 – 2 years post-infection. Cell populations expressing the ligand are shown on the x-axis (sending) and the cell populations expressing corresponding the receptor (receiving) are shown on the y-axis.

## Notes

### Competing Interest Statement

The authors have declared no competing interest.

